# Repeated replacement of an intrabacterial symbiont in the tripartite nested mealybug symbiosis

**DOI:** 10.1101/042267

**Authors:** Filip Husnik, John P. McCutcheon

## Abstract

Stable endosymbiosis of a bacterium into a host cell promotes cellular and genomic complexity. The mealybug *Planococcus citri* has two bacterial endosymbionts; remarkably, the gammaproteobacterium *Moranella endobia* lives in the cytoplasm of the betaproteobacterium *Tremblaya princeps*. These two bacteria, along with genes horizontally transferred from other bacteria to the *P. citri* genome, encode complementary gene sets that form a complex metabolic patchwork. Here we test the stability of this three-way symbiosis by sequencing host-symbiont genome pairs for five diverse mealybug species. We find marked fluidity over evolutionary time: while *Tremblaya* is the result of a single infection in the ancestor of mealybugs, the innermost gammaproteobacterial symbionts result from multiple replacements of inferred different ages from related but distinct bacterial lineages. Our data show that symbiont replacement can happen even in the most intricate symbiotic arrangements, and that pre-existing horizontally transferred genes can remain stable on genomes in the face of extensive symbiont turnover.

## Introduction

Many organisms require intracellular bacteria for survival. The oldest and most famous example are the eukaryotes, which depend on mitochondria (and, in photosynthetic eukaryotes, the chloroplasts or plastids) for the generation of biochemical energy (Gray and Doolittle 1982, Palmer 1997, Martin and Müller 1998, Embley and Martin 2006). But several more evolutionarily recent examples exist, where intracellular bacteria are involved in nutrient conversion or production from unbalanced host diets. For example, deep-sea tube worms, some protists, and many sap-feeding insects are completely dependent on intracellular bacteria for essential nutrient provisioning (Douglas 1989, Stewart et al. 2005, Nowack and Melkonian 2010). Some of these symbioses can form highly integrated organismal and genetic mosaics that in many ways resemble organelles (Nakayama and Ishida 2009, Husnik et al. 2013, Sloan et al. 2014, Luan et al. 2015). Like organelles, these endosymbionts have genomes encoding few genes (McCutcheon and Moran 2011, Moran and Bennett 2014), rely on gene products of bacterial origin that are encoded on the host genome (Nikoh et al. 2010, Nowack et al. 2011, Husnik et al. 2013, Sloan et al. 2014, Luan et al. 2015), and in some cases, import protein products encoded by these horizontally transferred genes back into the symbiont (Nowack and Grossman 2012, Nakabachi et al. 2014). The names given to these bacteria—endosymbiont, proto-organelle, or bona fide organelle—is a matter of debate (Theissen and Martin 2006, Keeling and Archibald 2008, McCutcheon and Keeling 2014, Keeling et al. 2015). What is not in doubt is that long-term interactions between hosts and essential bacteria have generated highly integrated and complex symbioses.

Establishment of a nutritional endosymbiosis is beneficial for a host by allowing access to previously inaccessible food sources. But strict dependence on intracellular bacteria can come with a cost: endosymbionts that stably associate with and provide essential functions to hosts often experience degenerative evolution (Sloan and Moran 2012a, Bennett and Moran 2013, Nakabachi et al. 2013, Manzano-Marín and Latorre 2014). This degenerative process is thought to be driven by long-term reductions in effective population size (N_e_) due to the combined effects of asexuality (loss of most recombination and lack of new DNA through horizontal gene transfer (HGT)) and host restriction (e.g. frequent population bottlenecks at transmission in vertically transmitted bacteria; Moran 1996). The outcomes of these processes are clearly reflected in the genomes of long-term endosymbionts: they are the smallest of any bacterium that is not an organelle, have the among the fastest rates of evolution measured for any bacterium (McCutcheon and Moran 2011, Moran and Bennett 2014), and are predicted to encode proteins and RNAs with decreased structural stability (Moran 1996, Fares et al. 2002a). In symbioses where the endosymbiont is required for normal host function, such as in the bacterial endosymbionts of sap-feeding insects, this degenerative process can trap the host in a symbiotic “rabbit hole,” where it depends completely on a symbiont which is slowly degenerating (Bennett and Moran 2015).

Unimpeded, the natural outcome of this degenerative process would seem to be extinction of the entire symbiosis. But extinction, if it does happen, is difficult to observe, and surely is not the only solution to dependency on a degenerating symbiont. For example, organelles are bacterial endosymbionts that have managed to survive for billions of years (Palmer 1997). Despite the reduced Ne of organelle genomes relative to the nuclear genome, eukaryotes are able to purge deleterious mutations that arise on organelle genomes, perhaps through a combination of host-level selection and the strong negative selective effects of substitutions on gene-dense organelle genomes (Popadin et al. 2013, Cooper et al. 2015). Extant organelle genomes also encode few genes relative to most bacteria, and it is also likely that a long history of moving genes to the nuclear genome has helped slow or stop organelle degeneration (Smith and Keeling 2015, Keeling et al. 2015). Some of the most degenerate insect endosymbionts also seem to have adopted a gene transfer strategy, although the number of transferred genes is far smaller compared to organelles. In aphids, mealybugs, psyllids, and whiteflies, some genes related to endosymbiont function are encoded on the nuclear genome, although in most cases these genes have been transferred from other bacteria, not the symbionts themselves (Nikoh et al. 2010, Husnik et al. 2013, Sloan et al. 2014, Luan et al. 2015). The other solution to avoid host extinction is to replace the degenerating symbiont with a fresh one, or to supplement it with a new partner. Examples of symbiont replacement and supplementation are replete in insects, occurring in at least the sap-feeding Auchenorrhyncha (McCutcheon and Moran 2007, Bennett and Moran 2013, Koga et al. 2013, Koga and Moran 2014), psyllids (Thao et al. 2000, Sloan and Moran 2012b), aphids (Lamelas et al. 2011, Vogel and Moran 2013, Manzano-Marín and Latorre 2014), lice (Smith et al. 2013), and weevils (Lefevre et al. 2004). When viewed over evolutionary time, it becomes clear that endosymbiosis can be dynamic—both genes and organisms come and go. It follows that any view of a symbiotic system established from just one or a few host lineages might provide only a snapshot of the complexity that built the observed relationship.

Mealybugs (Hemiptera: Cocoidea: Pseudococcidae) are a group of phloem sap-sucking insects that contain most of the symbiotic complexity described above. All of these insects depend on bacterial endosymbionts to provide them with essential amino acids missing from their diets, but this is accomplished in dramatically different ways in different mealybug lineages. One subfamily, the Phenacoccinae, have a single betaproteobacterial endosymbiont called *Tremblaya phenacola* which provides essential amino acids and vitamins to the host insect (Gruwell et al. 2010, Husnik et al. 2013). In the other subfamily of mealybugs, the Pseudococcinae, *Tremblaya* has been supplemented with a second bacterial endosymbiont, a gammaproteobacterium named *Moranella endobia* in the mealybug *Planococcus citri* (PCIT). While symbiont supplementation is not uncommon, what makes this symbiosis unique is its structure: *Moranella* stably resides in the cytoplasm of its partner bacterial symbiont, *Tremblaya princeps* (von Dohlen et al. 2001, Thao et al. 2002, Kono et al. 2008, McCutcheon and von Dohlen 2011).

The organisms in the nested three-way *P. citri* symbiosis are intimately tied together at the metabolic level. *Tremblaya princeps* PCIT (TPPCIT) has one of the smallest bacterial genomes ever reported, totaling 139 kb in length, encoding only 120 protein-coding genes, and lacking many translation-related genes commonly found in the most extremely reduced endosymbiont genomes (McCutcheon and von Dohlen 2011). Many metabolic genes missing in *Tremblaya* are present on the *Moranella endobia* PCIT (MEPCIT) genome; together with their host insect, these two symbionts are thought to work as a ‘metabolic patchwork’ to produce nutrients needed by all members of the consortium (McCutcheon and von Dohlen 2011). The symbiosis in *P. citri* is further supported by numerous horizontally transferred genes (HTGs) from several different bacterial donors to the insect genome, but not from *Tremblaya* or *Moranella*. These genes are up-regulated in the symbiotic tissue (bacteriome) and fill in many of the remaining metabolic gaps inferred from the bacterial endosymbiont genomes (Husnik et al. 2013).

Other data suggest additional complexity in the mealybug symbiosis. Phylogenetic analyses of the intra-*Tremblaya* endosymbionts show that while different lineages of mealybugs in the Pseudococcinae all possess gammaproteobacterial endosymbionts related to *Sodalis*, these bacteria do not show the co-evolutionary patterns typical of many long-term endosymbionts (Thao et al. 2002, Kono et al. 2008, López-Madrigal et al. 2014). These data raise the possibility that the innermost bacterium of this symbiosis may have resulted from separate acquisitions, or that the original intra-*Tremblaya* symbiont has been replaced in different mealybug lineages. What is not clear is when these acquisitions may have occurred and what effect they have had on the symbiosis. Here, using paired host-symbiont genome data from seven mealybug species (five newly generated for this study), we show that replacements of the intra-*Tremblaya* symbiont have likely occurred several times in mealybugs. We find that most nutrient-related bacterial horizontally transferred genes (HTGs) were acquired deep in the mealybug lineage, and that each subsequent symbiont supplementation and replacement have adapted to these pre-existing HTGs. Our data show that even complex and apparently highly integrated symbioses are subject to symbiont replacement, and that HTGs encoding parts of metabolic pathways can remain stable on host genomes in the face of endosymbiont turnover.

## Materials and Methods

### Symbiont genome sequencing, assembly, annotation, and analyses

Three mealybug species: *Maconellicoccus hirsutus* (pink hibiscus mealybug; MHIR; collection locality Egypt, Helwan), *Ferrisia virgata* (striped mealybug; FVIR; collection locality Egypt, Helwan), and *Paracoccus marginatus* (papaya mealybug; PMAR; collection locality Comoro Islands, Mayotte) were identified and provided by Thibaut Malausa (INRA, Sophia, France). *Trionymus perrisii* (TPER; collection locality Poland) samples were provided by Małgorzata Kalandyk-Kołodziejczyk (University of Silesia in Katowice, Poland). *Pseudococcus longispinus* samples (long-tailed mealybug; PLON) were collected by coauthor FH in a winter garden of Faculty of Science in Ceske Budejovice, Czech Republic. DNA vouchers and insect vouchers of adult females for slide mounting are available from FH. DNA was isolated from three to eight whole insects of all species by the Qiagen QIAamp DNA Micro Kit and each library was multiplexed on two thirds of an Illumina HiSeq 2000 lane and sequenced as 100 bp paired end reads. The *Maconellicoccus hirsutus* sample was sequenced on an entire MiSeq lane with v3 chemistry and 300 bp paired end mode. Both approaches generated sufficient coverage for both symbiont genomes and draft insect genomes. Adapter clipping and quality filtering was carried out in the Trimmomatic package (Bolger et al. 2014) using default settings. Read error correction (BayesHammer), de-novo assembly (k-mers K21, K33, K55, K77 for 100 bp data and K99, K127 for 300 bp data), and mismatch/short indel correction was performed by the SPAdes assembler v 3.5.0 (Bankevich et al. 2012). Additional endosymbiont-targeted long k-mer (91 and 241 bp) assemblies generated by the Ray v2.3.1 (Boisvert et al. 2010) and PRICE v1.2 (Ruby et al. 2013) assemblers were used to improve assemblies of complex endosymbiont regions.

Endosymbiont genomes were closed into circular-mapping molecules by combination of PCR and Sanger sequencing. General *Tremblaya* primers for closing of problematic regions such as the duplicated rRNA operon were designed to be applicable to most *Tremblaya princeps* species (Table S5). Given unclear GC skew in some of the species, the origin of replication was set to the same region as in already published *Tremblaya* and *Moranella* genomes to standardize comparative genomic analyses. Pilon v1.12 (Walker et al. 2014) and REAPR v1.0.17 (Hunt et al. 2013) were used to diagnose and improve potential misassemblies, collapsed repeats, and polymorphisms. Genome annotations and reannotations (*i.e.* for TPPCIT, MEPCIT and *Tremblaya phenacola* from *Phenacoccus avenae –* TPPAVE) were carried out by the Prokka v1.10 pipeline (Seemann 2014) with disabled default discarding of open reading frames (ORFs) overlapping tRNAs. Our new comparative data allowed us to re-annotate many genes and pseudogenes previously annotated as hypothetical proteins and to uncover pseudogene remnants (Table S1a). *Tremblaya* panproteome was curated manually with an extensive use of MetaPathways v2.0 (Konwar et al. 2013), PathwayTools v17.0 (Karp et al. 2010), and InterProscan v5.10 (Jones et al. 2014), and then used in Prokka as trusted proteins for annotation. This approach was used to obtain identical gene names for all seven *Tremblaya* genomes (TPPAVE, TPPCIT, TPMHIR, TPFVIR, TPPLON, TPPMAR, TPTPER; abbreviations combine TP with species abbreviations defined above). Transfer RNA and tmRNA regions were reannotated using tFind.pl wrapper [http://bioinformatics.sandia.gov/software]. *Tremblaya* pseudogenes were reannotated in the Artemis browser (Rutherford et al. 2000) based on genome alignment of all *Tremblaya* genomes.

Genomes of gammaproteobacterial symbionts were annotated as described for *Tremblaya* genomes, except that several approaches were employed to assist in pseudogene annotation. Proteins split into two or more open reading frames were joined into a single pseudogene feature. All proteins were then searched against the NCBI non-redundant protein database (NR) database and their length was compared. If the endosymbiont protein was shorter than 60% of its ten top hits, it was called a pseudogene unless it is known to be a bi-functional protein and at least one of its domains was intact. All intergenic regions were then screened by BlastX [e-value 1e^−4^] against NR to reveal pseudogene remnants.

Multi-gene matrices of conserved orthologous genes for Betaproteobacteria (49 genes) and Enterobacteriaceae (80 genes) were generated by the PhyloPhlAN package (Segata et al. 2013). Sequences of genes for 16S and 23S rRNA were downloaded from the NCBI nucleotide database and used for *Tremblaya* and *Sodalis*-allied species-rich phylogenies. All matrices were aligned by the MAFFT v6 L-INS-i algorithm (Katoh and Toh 2008). Ambiguously aligned positions were excluded by trimAL v1.2 (Capella-Gutiérrez et al. 2009) with the *–automated 1* flag set for likelihood-based phylogenetic methods. Maximum likelihood (ML) and Bayesian inference (BI) phylogenetic methods were applied to the single-gene and concatenated amino-acid alignments. ML trees were inferred using RAxML 8.2.4 (Stamatakis 2014) under the LG+G model with subtree pruning and re-grafting tree search algorithm (SPR) and 1000 bootstrap pseudo-replicates. BI analyses were conducted in MrBayes 3.2.2 (Ronquist and Huelsenbeck 2003) under the LG+I+G model with five million generations (*prset aamodel = fixed(lg), lset rates = invgamma ngammacat = 4, mcmcp checkpoint = yes ngen = 5000000*). Concatenated 16S-23S rRNA gene phylogenies for mealybug endosymbionts were inferred as above, except that the GTR+I+G model was used. For BI analyses, a proportion of invariable sites (I) was estimated from the data and heterogeneity of evolutionary rates was modeled by the four substitution rate categories of the gamma (G) distribution with the gamma shape parameter (alpha) estimated from the data. Exploration of MCMC convergence and burn-in determination was performed in AWTY [http://ceb.csit.fsu.edu/awty] and Tracer v1.5 [http://evolve.zoo.ox.ac.uk]. Additionally, concatenated protein and Dayhoff6 recoded datasets were analyzed under the CAT+GTR+G model in PhyloBayes MPI 1.5a (Lartillot et al. 2013). Posterior distributions obtained under four independent PhyloBayes runs were compared using tracecomp and bpcomp programs and runs were considered converged at maximum discrepancy value < 0.1 and minimum effective size > 100.

*Tremblaya* genomes were aligned using progressiveMauve v2.3.1 (Darling et al. 2010). Clusters of orthologous genes were generated using OrhoMCL v1.4 (Li et al. 2003). Orthologs missed due to low homology (BLAST e-value 1e^−5^) were curated with help of identical gene order and annotations. All genomes were visualized as linear with links connecting positions of orthologous genes in Processing3 [https://processing.org/]. Additional figures were drawn or curated in Inkscape [https://inkscape.org/].

### Contamination screening and filtering of draft mealybug genomes

The presence of additional species such as facultative symbionts, environmental bacteria, and contamination in the genome data was visualized by the Taxon-annotated GC-Coverage plots (TAGC; http://drl.github.io/blobtools/; Kumar et al. 2013, Koutsovoulos et al. 2015) and the tool was also used to extract contigs of two gammaproteobacterial symbionts from the *P. longispinus* mealybug and *Wolbachia* sp. from the *M. hirsutus* mealybug. We confirmed that there were no other organisms present in our data at high coverage except the expected endosymbionts. Although there are now reliable methodologies to remove majority of contamination from data sequenced using several independent libraries (Koutsovoulos et al. 2015, Delmont and Eren 2016), recognizing low-coverage contamination (in our case mostly of bacterial, human, and plant origin) from single library sequencing data can be problematic. Using the TAGC tool, we were able to recognize low-coverage *Propionibacterium* spp. and human contamination in several of the samples (megablast e-value 1e^−25^), and plant contamination in the *P. longispinus* sample. These short sequences were filtered out and also all (non-symbiont) contigs or scaffolds shorter than 200 bp and/or having coverage lower than 3x were excluded from the total assemblies.

### Draft insect genomes and horizontal gene transfers

Endosymbiont contigs and PhiX contigs (from the spike-in of Illumina libraries) were excluded from assemblies and insect genome assemblies were evaluated by the Quast v.2.3 tool (Gurevich et al. 2013) for basic assembly statistics and by the CEGMA v2.5 (Parra et al. 2007) and BUSCO v1.1 (Simão et al. 2015) with Arthropoda dataset for gene completeness (Table S2). Lacking RNA-Seq data to properly annotate the draft genomes, only preliminary gene predictions were carried out by unsupervised GeneMark-ES (Lomsadze et al. 2005) runs to get exon structures for scaffolds with HGTs.

Horizontally transferred genes previously identified in the *P. citri* genome were used as queries for BlastN, tBlastN, and tBlastX searches against custom databases made of scaffolds from individual species. Additionally, two approaches were used to minimize false negative results possibly caused by highly diverged and/or fragmented HGTs undetected by Blast searches. First, nucleotide alignments of individual HGTs (see above) were used as HMM profiles in nhmmer (Wheeler and Eddy 2013) searches against scaffolds of individual assemblies. Second, Blast databases were made out of all raw fastq reads and searched by tBlastN using protein HGTs from *P. citri* as queries.

Lineage-specific candidates of horizontal gene transfer were detected as reported previously (Husnik et al. 2013) using NR database (downloaded March 17, 2015). We used stringent screening criteria: only genes present on long scaffolds containing insect genes or present in several mealybug genomes were considered as strongly supported HGT candidates here (Table S4). Moreover, all scaffolds of HGT candidates presented here were confirmed by mapping raw read data and manually examined for low-coverage regions and potential mis-assemblies created by the joining of low-coverage contigs of bacterial contaminants with bona-fide insect contigs.

A multi-gene mealybug phylogeny was inferred as above using 419 concatenated protein sequences of the core eukaryotic proteins identified from the six mealybug genomes by the CEGMA package. Phylogenetic trees for individual HGTs were inferred as reported previously (Husnik et al. 2013) except that the workflow was implemented using ETE3 Python toolkit (Huerta-Cepas et al. 2010).

## Results

### Overview of our sequencing efforts

We generated genome data for five diverse Pseudococcinae mealybug species, in total closing nine symbiont genomes into single circular-mapping molecules (five genomes from *Tremblaya*; four from the *Sodalis*-allied gammaproteobacterial symbionts) (Table 1). Unexpectedly, we detected gammaproteobacterial symbionts in *Ferrisia* and *Maconellicoccus* species, which were not previously reported to harbor intrabacterial symbionts inside *Tremblaya* cells (Figure 1,2,3). We also found that *P. longispinus* harbored two gammaproteobacterial symbionts, each with complex genomes larger than 4Mbp; these were left as draft assemblies of 231 contigs with total assembly size of 8,191,698 bp and N50 of 82,6 kbp (Table 1).

**Figure 1.**
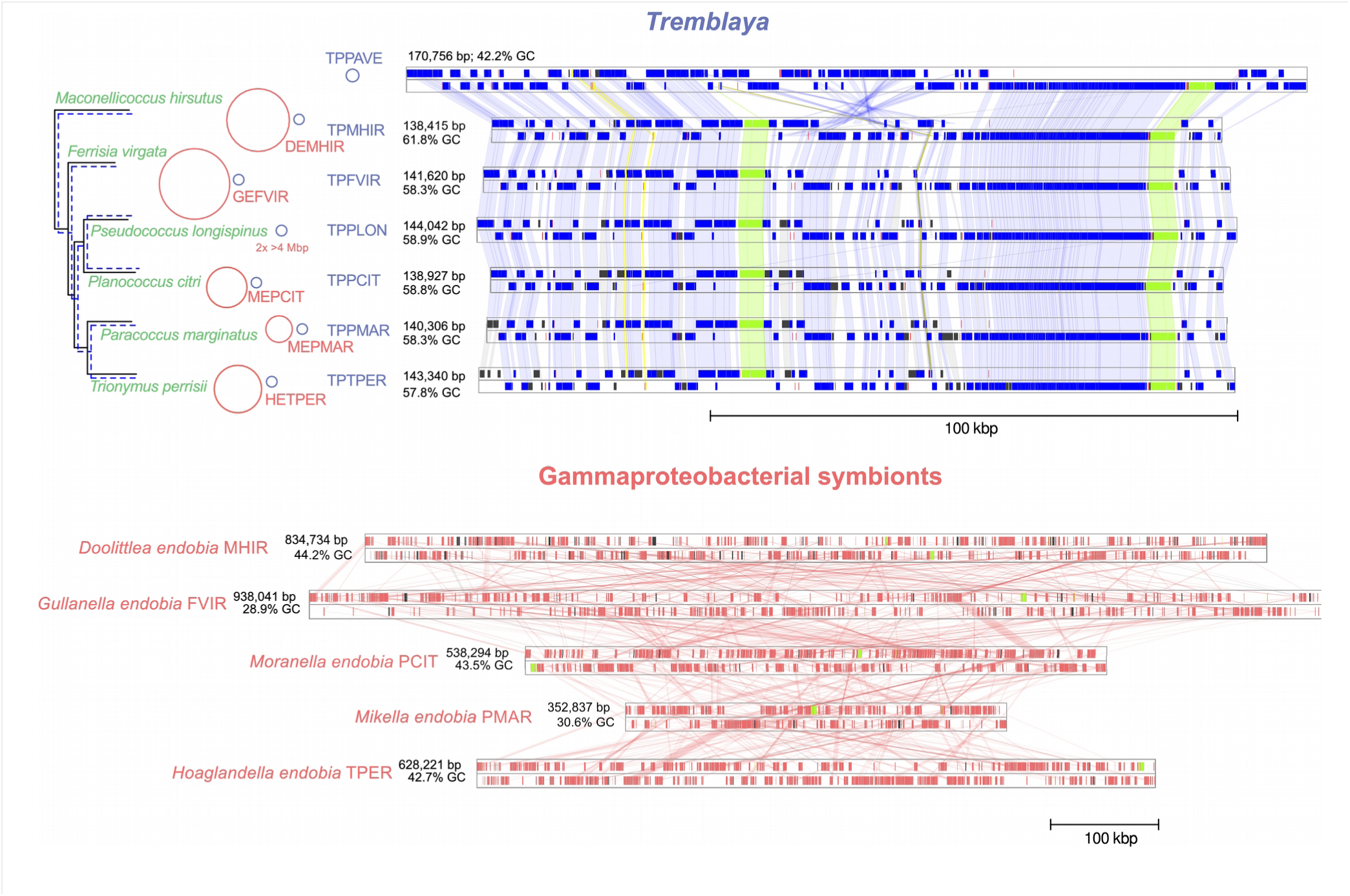
Genome size and structure of the mealybug endosymbionts. Linear genome alignments of seven *Tremblaya* genomes (top, blue) are contrasted with linear genome alignments of five genomes of their respective gammaproteobacterial symbionts (bottom, red). The *Tremblaya* genomes are perfectly co-linear and similar in size, while the gammaproteobacterial genomes are highly rearranged and different in size. Alignments are ordered based on schematic mealybug-*Tremblaya* phylogeny (original phylogenies in Figure S1) and accompanied by basic genome statistics (detailed genome statistics in Table 1). Gene boxes are colored according to their category: proteins in blue, pseudogenes in grey, ribosomal RNAs in green, non-coding RNAs in yellow, and transfer RNAs in red.

**Figure 2.**
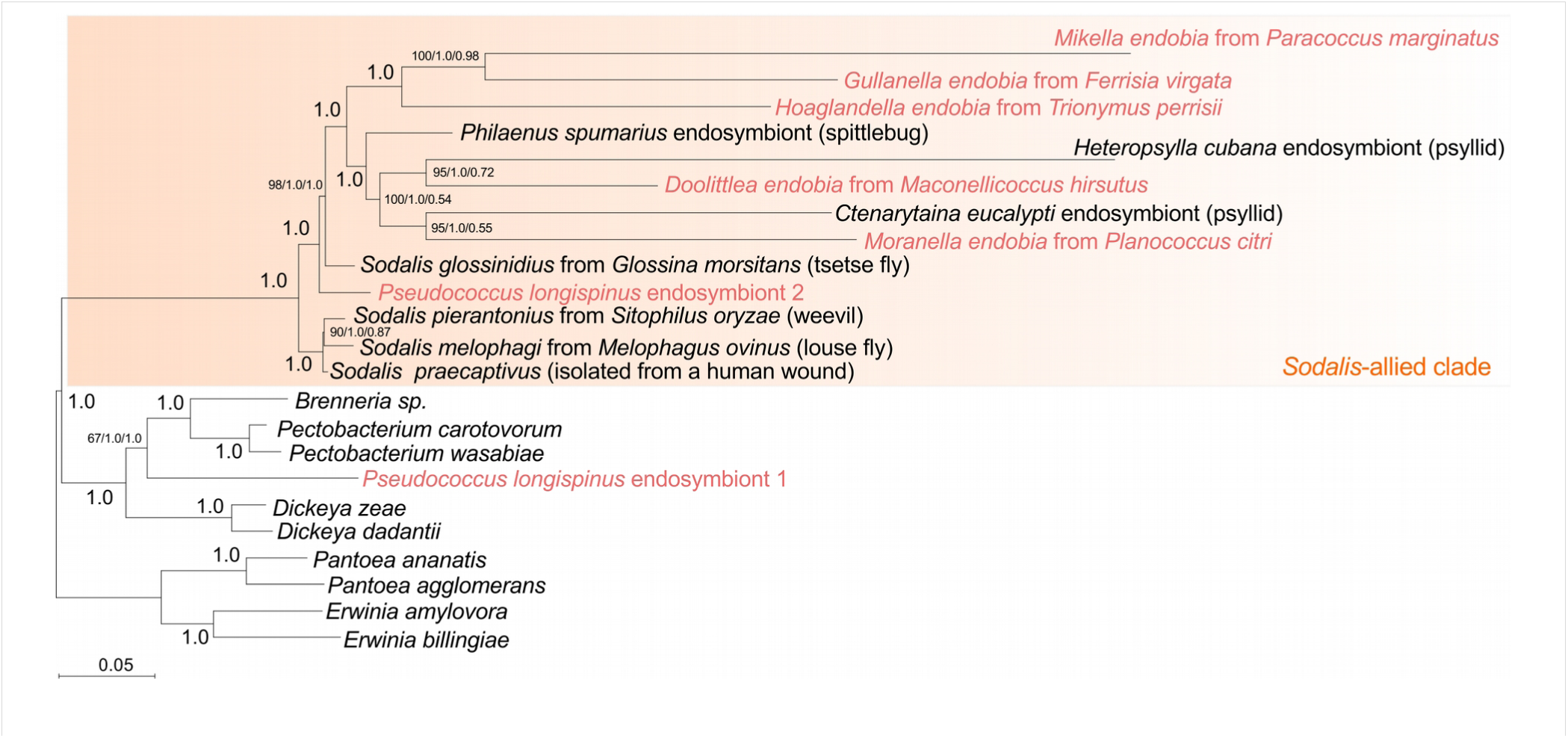
The intrabacterial mealybug symbionts are members of the *Sodalis* clade of gammaproteobacteria. A multi-gene phylogeny of *Sodalis*-allied insect endosymbionts and closely related Enterobacteriaceae (Gammaproteobacteria) was inferred from 80 concatenated proteins under the LG+G evolutionary model in RaxML v8.2.4. Mealybug endosymbionts are highlighted in red. Values at nodes represent bootstrap pseudoreplicates from the ML analysis, posterior probabilities from BI topology inferred under LG+I+G model, and posterior probabilities from BI topology inferred from Dayhoff6 recoded dataset under the CAT+GTR+G model in PhyloBayes.

**Figure 3.**
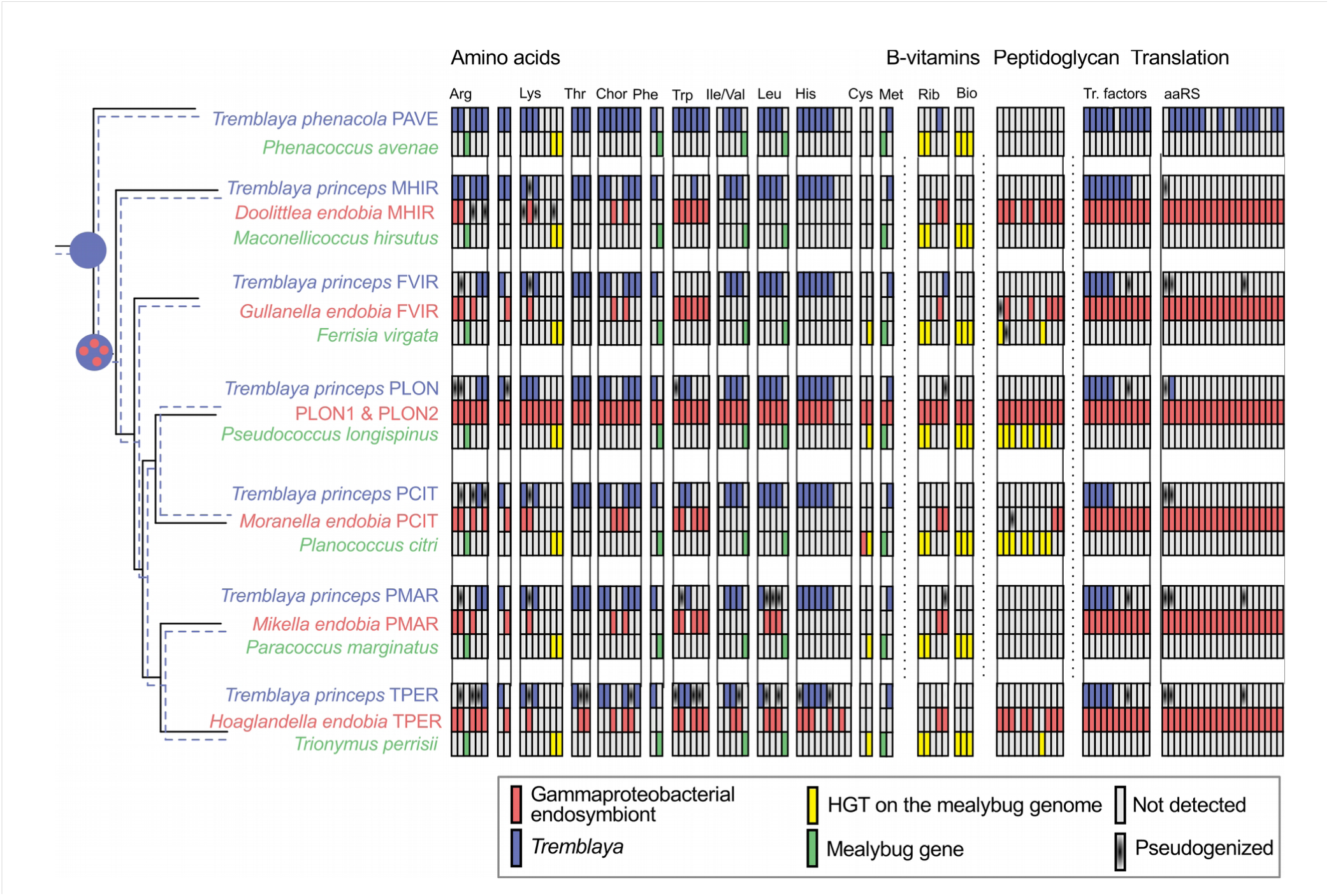
A complex history of gene retention, loss, and acquisition in the mealybug symbiosis. Retention of selected biosynthetic pathways (amino acids, B-vitamins, and peptidoglycan), translation-related genes, and horizontally transferred genes visualized by colored rectangles for seven mealybug symbiotic systems. For each mealybug species, row one represents *Tremblaya* (in blue), row two represents its gammaproteobacterial symbionts (in red), and row three represents the host genome (insect genes in green, HGTs in yellow). Missing genes are shown in grey and recognizable pseudogenes are shown with black radial gradient. Raw data used for this table (including gene names) are available in the Tab S2.

**Tab 1.**
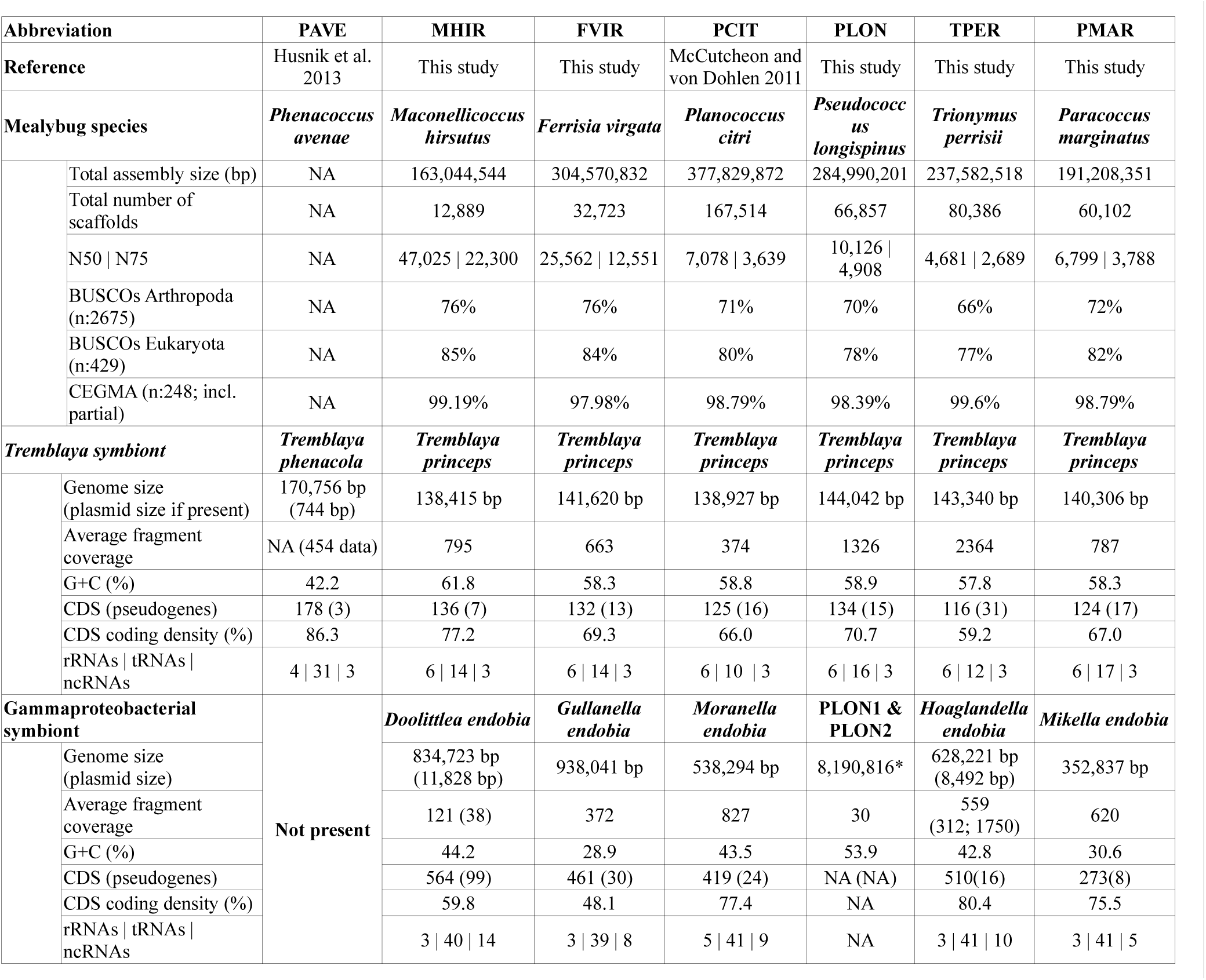
Genome statistics for mealybug endosymbionts and draft mealybug genomes. The asterisk denotes combined assembly size for both gammaproteobacterial symbionts in PLON. *Hoaglandella endobia* codes two plasmids of 3,244 and 5,248 base pairs. Extended assembly metrics for draft mealybug genomes are available as Table S2.

We also assembled five mealybug draft genomes (Table 1). Because our assemblies were generated only from short-insert paired-end data, the insect draft genomes consisted primarily of numerous short scaffolds (Table S2).

### *Tremblaya* genomes are stable in size and structure, the gammaproteobacterial genomes are not

All five *T. princeps* genomes (those that have a gammaproteobacterial symbiont) are completely syntenic with each other and similar in size, ranging from 138 kb to 143 kb (Figure 1). The gene contents are also similar, with 107 protein-coding genes shared in all five *Tremblaya* genomes. All differences in gene content come from gene loss or nonfunctionalization in different lineages (Figure 1). Four pseudogenes (*argS*, *mnmG*, *lpd*, *rsmH*) are shared in all five *T. princeps* genomes, indicating that some pseudogenes are retained in *Tremblaya* for long periods of time. Pseudogene numbers were notably higher, and coding densities lower, in *P. marginatus* and *T. perrisii* (Figure 1 and Table 1).

In contrast to the genomic stability observed in *Tremblaya*, the genomes of the gammaproteobacterial symbionts vary dramatically in size, coding density, and gene order (Figures 1 and 3; Table 1). These genomes range in size from 353 to 8,000 kb (from the presence of two ~4,000 kb genomes in *P. longispinus*), and are all notably different from the 539 kb *Moranella* genome of *P. citri* (McCutcheon and von Dohlen 2011).

### Phylogenetic analyses confirm the intra-*Tremblaya* gammaproteobacterial symbionts result from multiple infections

The lack of conservation in gammaproteobacterial genome size and structure, combined with data showing their phylogeny does not mirror that of their mealybug or *Tremblaya* hosts (Thao et al. 2002, Kono et al. 2008; see also Figure S1), supports early hypotheses that the gammaproteobacterial symbionts of diverse mealybug lineages result from multiple unrelated origins (Thao et al. 2002, Kono et al. 2008). Although the *Sodalis*-allied clade is extremely hard to resolve due to low taxon-sampling of facultative and free-living relatives, nucleotide bias, and rapid evolution in obligate symbionts, none of our analyses indicate a monophyletic group of mealybug symbionts congruent with the host and *Tremblaya* trees (Figures 2 and S1).

### Draft insect genomes reveal the timing of mealybug horizontal gene transfers

Gene annotation of low-quality draft genome assemblies is known to be problematic (Denton et al. 2014). We therefore verified that our mealybug assemblies were sufficient for our purpose of establishing gene presence or absence by comparing our gene sets to databases containing core eukaryotic (CEGMA) and Arthropod (BUSCO) gene sets. CEGMA scores surpass 98% in all of our assemblies, and BUSCO Arthropoda scores range from 66 to 76% (Table S2). We note that the low scores against the BUSCO database likely reflect the hemipteran origin of mealybugs rather than our fragmented assembly; the high-quality pea aphid genome (International Aphid Genomics Consortium 2010) scores 72% using identical settings. We thus conclude that our mealybug draft assemblies are sufficient for determining the presence or absence of bacterial HGTs.

We first sought to confirm that HTGs found previously in the *P. citri* genome (Husnik et al. 2013) were present in other mealybug species (Tables S3 and S4), and to establish the timing of these transfers. (Consistent with our previous findings (Husnik et al. 2013), there were no well-supported HGTs of *Tremblaya* origin detected in any of our mealybug assemblies.) Our data show that the acquisition of some HTGs (*bioABD*, *ribAD*, *dapF*, *lysA*, *tms*, AAA-atpases) predated the Phenacoccinae/Pseudococcinae divergence, and thus the acquisition of the gammaproteobacterial endosymbiont (Figure 3). These old HGTs mostly involve amino acid and B-vitamin metabolism, are usually found on longer insect scaffolds which contain several essential insect genes, and are syntenic across mealybug species (Figure 4). In each of these cases, no other bacterial genes or pseudogenes were found within the scaffolds (Tables S3 and S4), suggesting that these HTGs resulted from the transfer of small DNA fragments, or that flanking bacterial DNA from larger transfers was lost soon after the transfer was established. The origin of some of these transfers (*bioAB*) predate the entire mealybug lineage, since they are found in the genome of the whitefly *Bemisia tabaci* (Luan et al. 2015).

**Figure 4.**
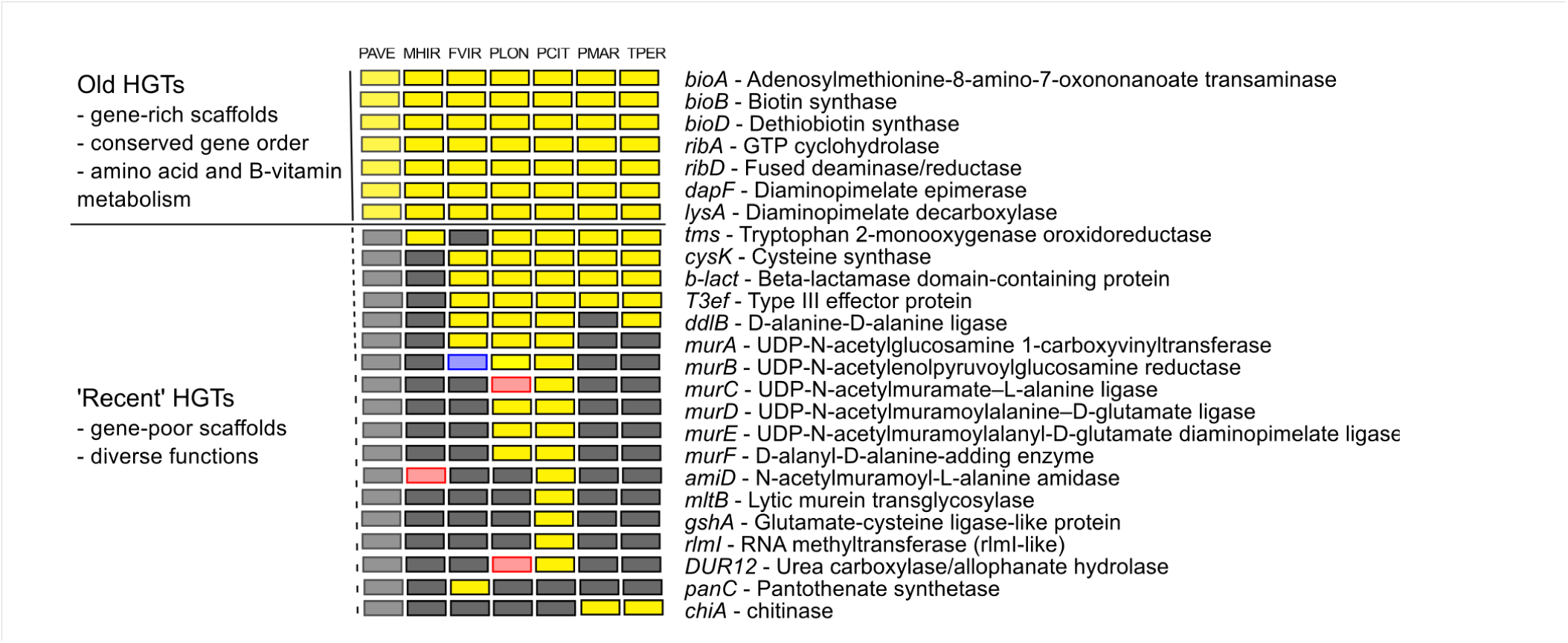
HGTs detected in individual mealybug species. Retention of horizontal gene transfer candidates detected across all mealybug species (yellow: gene present, grey: gene not detected, red: different phylogenetic origin, blue: possible pseudogene).

We find that several HTGs were acquired after the divergence of the *Maconellicoccus* clade (*cysK, b-lact, T3ef, ddlB*). One of these genes, cysteine synthase A (*cysK*), clusters with sequences from other *Sodalis*-allied bacteria, consistent with a possible origin from an early gammaproteobacterial intrabacterial symbiont (Figure S3f). We note that *cysK* has undergone tandem duplication in *P. longispinus*, *F. virgata*, and *P. citri* (Tables S3 and S4), which was also observed for several other HTGs (*tms*, *b-lact*, *T3ef, chiA*, ankyrin repeat proteins and AAA ATPases). Most of the HTGs detected from only one or two mealybug species are related to peptidoglycan metabolism and were assembled on shorter scaffolds with very few insect genes on them. Possible HGT losses of *tms* in *F. virgata* and *ddlB* in *P. marginatus* were detected based on our assemblies. Except in three cases (*amiD*, *murC*, and DUR1), identical HGT candidates detected from several mealybug species shared significant amount of sequence similarity and clustered as a single clade in our phylogenies (Figure S3a-u), suggesting that these transfers results from single events.

### Evolution of the metabolic patchwork

We first found complementary patterns of gene loss and retention between *Tremblaya*, *Moranella*, and the mealybug host in the *P. citri* symbiosis (McCutcheon and von Dohlen 2011, Husnik et al. 2013). Our new comparative genomic data allow us to see how genes are retained or lost in different genomes in multiple lineages that have gammaproteobacterial symbionts of different inferred ages (Figure 3). These new data also allow us to observe how new symbionts evolve in response to the presence of both pre-existing symbionts and horizontally transferred genes.

Our analysis is simplified by the availability of the *Tremblaya* genome from PAVE, which lacks a gammaproteobacterial endosymbiont (Husnik et al. 2013). In all other species, our assumption is that the gene set of *Tremblaya* PAVE was present at the acquisition of the first gammaproteobacterial symbiont, and that any gene loss seen in *Tremblaya* lineages is in response to this event. Overall our data point to an extremely complex pattern of gene loss and retention in the mealybug symbiosis (Figure 3). Some pathways, such as those for the production of lysine, phenylalanine, and methionine, show the stereotypical patchwork pattern in all mealybugs, with gene retention is interspersed between *Tremblaya* and its gammaproteobacterial endosymbiont. Gene retention patterns from many other pathways, though, show much less predictable patterns. The isoleucine, valine, leucine, threonine, and histidine pathways show a strong tendency towards *Tremblaya*-dominated biosynthesis in *M. hirsutus*, *F. virgata*, and *P. citri* (that is, retention in *Tremblaya* and loss in the gammaproteobacterial symbiont), but with a clear shift towards gammaproteobacterial-dominated biosynthesis in *P. marginatus* and *T. perrisii*. Other pathways, such as tryptophan, show gammaproteobacterial dominance in all mealybug symbioses, but with reliance on at least one *Tremblaya* gene in *P. citri*, *P. marginatus*, and *T. perrisii*. In the arginine pathway, gene retention is dominated by *Tremblaya* in *M. hirsutus*, but by the gammaproteobacterial endosymbiont in all other lineages, with sporadic loss of *Tremblaya* genes in different lineages. Overall, *M. hirsutus* encodes the most *Tremblaya* genes and the fewest gammaproteobacterial genes, while *T. perrisii* shows the opposite pattern. Interestingly, these patterns between mealybug lineages do not strongly correspond to gammaproteobacterial genome size (Table 1 and Figure 3).

### Gene retention patterns for translation-related genes in *Tremblaya*

In contrast to metabolic genes involved in nutrient production, the retention patterns for genes involved in translation vary little between mealybug species (Figure 3). As first shown in *Tremblaya* PCIT (McCutcheon and von Dohlen 2011), none of the additional *Tremblaya* genomes we report here encode any functional amino-acyl tRNA synthetase (aaRS) with an exception of one likely functional gene (*cysS*) in *T. princeps* PLON, which is present as a pseudogene in several other lineages of Tremblaya. Further, all *Tremblaya* genomes have lost key translational control proteins that are typically retained even in the smallest endosymbiont genomes, such as ribosome recycling factor (*rrƒ*), L-methionyl-tRNA^fMet^ N-formyltransferase (*ƒmt*), and peptide deformylase (*deƒ*). The translational release factors RF-1 and RF-2 (*prƒAB*) and elongation factor EF-Ts (*tsƒ*) are present only in the gene-rich *T. princeps* MHIR genome, but absent or pseudogenized in all other *T. princeps* genomes. Initiation factors IF-1, IF-2, and IF-3 (*inƒABC*), and elongation factors EF-Tu and EF-G (*tuƒA* and *ƒusA*) are retained in all *Tremblaya* genomes, as are most ribosomal proteins (Table S1).

### Taxonomy of mealybug endosymbionts

The naming convention in the field of insect endosymbiosis has been to keep the species names constant for lineages of endosymbiotic bacteria, even if they exist in different species of host insects. The host is denoted by appending a specific abbreviation to the end of the endosymbiont name (e.g., *Tremblaya princeps* PCIT for *T. princeps* from *Planoccocus citri*). However, our data show that the intra-*Tremblaya* gammaproteobacterial symbionts are not from the same lineage; they result from independent endosymbiotic events from clearly discrete lineages within the *Sodalis* clade (Figure 2). Following convention, we have chosen to give these gammaproteobacteria different genus names, but to unite them by retaining the ‘*endobia*’ species denomination for each one, as in *Moranella endobia*).

We propose the following Candidatus status names for the four lineages of intra-*Tremblaya* gammaproteobacterial symbionts of mealybugs for which we have completed a genome. First, *Candidatus* Doolittlea endobia MHIR for the endosymbiont from *Maconellicoccus hirsutus*. This name honors the American evolutionary biologist W. Ford Doolittle (1941-) for his contributions to our understanding of HGT and endosymbiosis. *Candidatus* Gullanella endobia FVIR for the endosymbiont from *Ferrisia virgata*. This name honors the Australian entomologist Penny J. Gullan (1952-) for her contributions to numerous aspects of mealybug biology and taxonomy. *Candidatus* Mikella endobia PMAR for the endosymbiont from *Paracoccus marginatus*. This name honors the Canadian biochemist Michael W. Gray (1943-) for his contributions to our understanding of organelle evolution. *Candidatus* Hoaglandella endobia TPER for the endosymbiont from *Trionymus perrisii*. This name honors the American biochemist Mahlon B. Hoagland (1921-2009) for his contributions to our understanding of the genetic code, including the co-discovery of tRNA. All of the names we propose could be extendible to related mealybugs species (e.g. *Gullanella endobia* for other members of the *Ferrisia* clade) if future phylogenetic analyses show that these symbionts are part of the same lineage. For simplicity, we use all endosymbiont names without the *Candidatus* denomination.

## Discussion

### Diversity of intrabacterial symbiont genomes suggests multiple replacements

Phylogenetic analyses based on rRNA and protein-coding genes from the gammaproteobacterial endosymbionts of mealybugs first indicated their origins from multiple unrelated bacteria (Thao et al. 2002, Kono et al. 2008). What was less clear from these data was the order and timing of the gammaproteobacterial infections, and how these infections affected the other members of the symbiosis. We imagine three possible scenarios that could explain our phylogenetic and genomic data (Figure 5). The first is that there was a single gammaproteobacterial acquisition in the ancestor of the Pseudococcinae which has evolved idiosyncratically as mealybugs diversified over time, leading to seemingly unrelated genome structures and coding capacities (the “idosyncratic” scenario; Figure 5A). The second is that the gammaproteobacterial infections occurred independently, each establishing symbioses inside *Tremblaya* in completely unrelated and separate events (the “independent” scenario; Figure 5B). The third is that there was a single gammaproteobacterial acquisition in the Pseudococcinae ancestor that has been replaced in some mealybug lineages over time (the “replacement” scenario; Figure 5C). The idosyncratic scenario is easy to disregard, because while acquisition of a symbiont followed by rapid diversification of the host might result in different patterns of genome evolution in different lineages, it should result in monophyletic clustering in phylogenetic trees. Previous phylogenetic work, as well as our phylogenomic data (Figure 2) show that the gammaproteobacteria that have infected different mealybugs have originated from clearly distinct (and well-supported) bacterial lineages.

**Figure 5.**
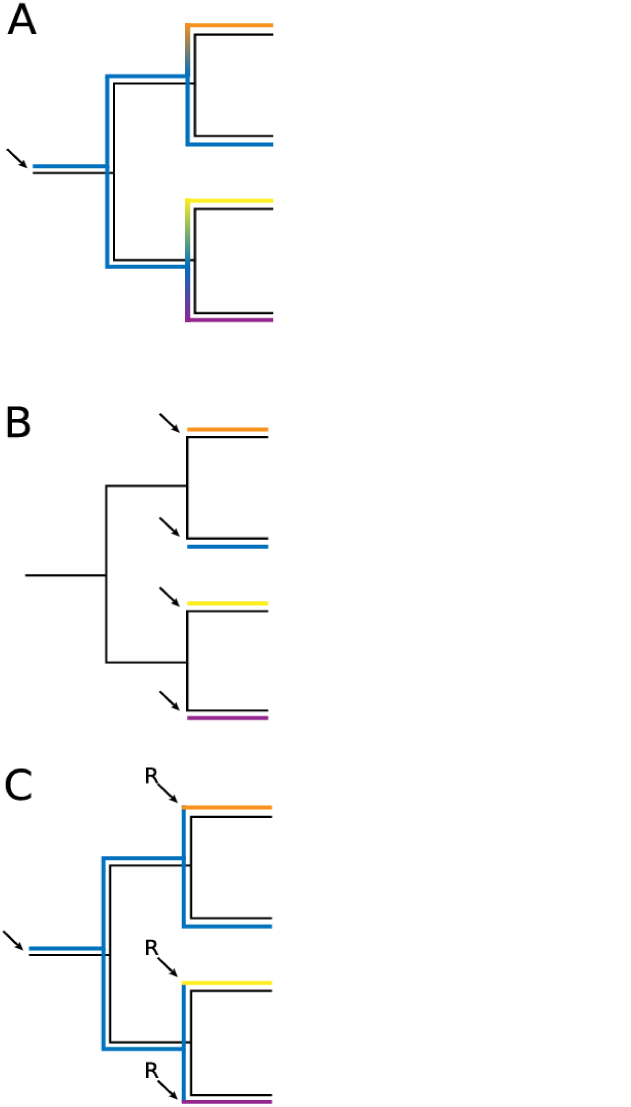
Three possible scenarios that built the mealybug symbiosis. Independent gammaproteobacterial acquisitions are shown as arrows, replacements are noted with and “R” above the arrow. Colors represent the different gammaproteobacterial genomes shown in Figure 1. (A) The idiosyncratic scenario, where a single gammaproteobacterial acquisition evolved differently as mealybugs diverged, leading to different genome sizes and structures in extant mealybugs. (B) The independent scenario, where the different sizes and structures of the gammaproteobacterial genomes shown in Figure 1 result from completely independent acquisitions. (C) The replacement scenario, where the different sizes and structures of the gammaproteobacterial genomes shown in Figure 1 result from several replacements of an ancestral gammaproteobacterial symbiont.

The independent and replacement scenarios are more difficult to tell apart with our data, but we favor the replacement model, primarily because of the large differences in size we observe in the gammaproteobacterial genomes. Genome size is strongly correlated to endosymbiotic age in bacteria, especially at the onset of symbiosis, when genome reduction can be rapid (Moran and Mira 2001, Frank et al. 2002, Moran 2002, Moran et al. 2008, Moya et al. 2008). Most relevant to our argument here is the speed with which genome reduction has been shown to take place in *Sodalis*-allied bacteria closely related to the gammaproteobacterial symbionts of mealybugs (Clayton et al. 2012, Oakeson et al. 2014, Koga and Moran 2014). It has been estimated that as much as 55% of an ancestral genome can be lost upon the transition to endosymbiosis in a mere ~28,000 years in one *Sodalis* lineage, barely enough time for 1% sequence divergence to accumulate between the new symbiont and a free-living relative (Clayton et al. 2012). Our general assumption is therefore that recently established endosymbionts should have larger genomes than older symbionts. However, we note that genome reduction is not a deterministic process related to time, especially as the symbiosis ages. It is clear that in some insects housing pairs of ancient symbionts with highly reduced genomes, the older endosymbiont can have a larger genome than the newer symbiont (McCutcheon and Moran 2010).

The evidence for recent replacement is most obvious in *P. longispinus* (Table 1 and Figure 3). This symbiosis harbors two related gammaproteobacterial symbionts (Rosenblueth et al. 2012), each with a rod-like cell shape, although it is currently unclear if both bacteria reside within *Tremblaya* (Gatehouse et al. 2012). Both of these genomes are about 4 Mb in size (this study), approximately the same size as the recently acquired *Sodalis* symbionts from tsetse fly (4.3 Mb; Toh et al. 2006, Belda et al. 2010) and the rice weevil (4.5 Mb; Oakeson et al. 2014). These morphological and genomic features, as well as their short branches in Figure 2, all suggest the *P. longispinus* gammaproteobacterial symbionts are recent acquisitions in the *P. longispinus* symbiosis. The *P. longispinus* replacement seems so recent that the stereotypical complementary patterns of gene loss and retention have not had time to accumulate between the gammaproteobactia and *Tremblaya* (Figure 3). However, *Tremblaya* PLON is missing the same translation-realted genes (aside from *cysS*) as all other *Tremblaya*, indicating that it has long ago adapted to the presence of a (now eliminated) bacterium living in its cytoplasm. Comprehensive analyses of the two gammaproteobacterial genomes from *P. longispinus* are on-going and will be published elsewhere.

We thus hypothesize that the larger, gene-rich gammaproteobacterial genomes we describe here are the result of symbiont replacements of an ancestral gammaproteobacterial endosymbiont rather than completely independent infections in different mealybug lineages. We suspect that the massive loss in key translation-related genes (Figure 3) occurred in response to the first gammaproteobacterial infection, which subsequently required all subsequent replacement events to also reside within the *Trembalay* cytoplasm. We also note that in at least one other case bacteria from the *Sodalis* group have established multiple repeated infections in a replacement-like pattern (Smith et al. 2013). It is tempting to speculate that the 353 kb *Mikella* PMAR genome is the ancestral intra-*Tremblaya* symbiont lineage that has not been replaced, or at least has not been recently replaced. However, because the relevant clades split right after the Phenacoccinae/Pseudococcinae divergence—that is, right at the acquisition of the first gammaproteobacterial symbiont—much richer taxon sampling would be needed to test the hypothesis that this was in fact the original symbiont lineage (Figure 2).

### How did the bacteria-within-a-bacterium structure start, and why does it persist?

In extreme cases of endosymbiotic genome reduction, genes required for the generation of a cell envelope, among other fundamental processes, are lost (McCutcheon and Moran 2011, Moran and Bennett 2014). This is true for *Tremblaya*, where even the largest genome (from *P. avenae*, which lacks a gammaproteobacterial symbiont) encodes no genes for the production of fatty acids or peptidoglycan (Husnik et al. 2013). Therefore, when the first gammaproteobacterial endosymbiont established residence in *Tremblaya*, it likely invaded a membrane system that was more eukaryotic than bacterial in nature. We therefore consider the possibility that the symbiosis first started from bacterivory of a gammaproteobacterium by *Tremblaya* unlikely. It is now clear that bacteria in the *Sodalis* group are very good at establishing intracellular infections in insects (Dale et al. 2001, Smith et al. 2013, Hosokawa et al. 2015); it now also seems that *Sodalis* are good at establishing intracellular infections inside *Tremblaya*. But why did it start in *Tremblaya*? We can think of two related possibilities. The first is that it was easier to use the established transport system between the insect cell and *Tremblaya* (Duncan et al. 2014) than to evolve a new one. The second is that the insect immune system likely does not target *Tremblaya* cells, so it is an ideal hiding place for facultative symbiont, at least at the beginning of the intrabacterial symbiosis. After the loss of critical translation-related genes, the symbiosis would persist as bacteria-within-a-bacterium because no other structure is possible. We note that *Sodalis* and *Arsenophonus*-allied symbionts were recently suggested to sometimes reside within *Sulcia* cells in the leafhoppers *Cicadella viridis* and *Macrosteles laevis* (Michalik et al. 2014, Kobiałka et al. 2015). Although these studies were based only on electron microscopy imaging and not confirmed by specific probes (e.g. with FISH), it is possible that intrabacterial symbiosis is not a rare event in insect systems.

### Evolution of organelles and the timing of HGT

The parallels between bona fide organelles and obligate beneficial endosymbionts are clear: both have undergone large levels of gene loss and genome reduction (Moran and Bennett 2014, Smith and Keeling 2015), the genomes of both are often (Tamas et al. 2002, Burger et al. 2013)—but not always (Sloan et al. 2012, Campbell et al. 2015) — stable over long periods of time, both rely on horizontally transferred genes from bacteria to the host genome (Timmis et al. 2004, Nowack and Grossman 2012, Nakabachi et al. 2014), and both are required for host survival. But are the beneficial endosymbionts of insects, protists, and other eukaryotes really comparable to mitochondria and plastids? In some sense they are not, because nothing in biology is or can be: The cellular organelles each evolved once and are fundamental to life as we know it. But outside of age and perceived specialness (Booth and Doolittle 2015), many of the mechanistic and evolutionary outcomes of intimate endosymbiosis seem similar between organelles and insect endosymbionts (McCutcheon and Keeling 2014). We argue that these other, younger symbioses may tell us something about how the mitochondria and plastids came to be, at the very least by revealing what types of evolutionary events are possible as stable intracellular relationships proceed along the path of integration.

It is widely accepted that the mitochondria found across eukaryotes are related back to a single common alphaproteobacterial ancestor (Wang and Wu 2015), and that the plastids result from a single cyanobacterial infection (Ochoa de Alda et al. 2014). What is less clear is what happened before these endosymbiont lineage were fixed into organelles. The textbook concept is that a bacterium was taken up by a host cell, transferred most of its genes, and became the mitochondrion or plastid (Booth and Doolittle 2015). This idea becomes more complicated when the taxonomic affiliation of bacterial genes on eukaryotic genomes are examined (Kurland and Andersson 2000, Zimorski et al. 2014, Ku et al. 2015a, Gray 2015). For example, only about 20% of mitochondria-related horizontally transferred genes have strong alphaproteobacterial phylogenetic affinities (Gray 2015). The signals for the remaining 80% are either too weak to confidently place the gene, or show clear affiliation with other bacterial groups (Kurland and Andersson 2000, Gray 2015). Hypotheses that explain these data fall roughly into two camps. Some imagine a gradual process where multiple taxonomically diverse endosymbioses may have occurred— and transferred genes—before the final alphaproteobacterial symbiont was fixed. That is, the mitochondria arrived rather late in the evolution of a cell that already contained many bacterial genes resulting from HGT of previous symbionts (Larkum et al. 2007, Ettema 2016, Pittis and Gabaldón 2016). Others favor a more abrupt ‘mitochondria early’ scenario, where an endosymbiont with a taxonomically diverse mosaic genome made the transition to becoming the mitochondrion in a single endosymbiotic event, transferring its genes during the process. In this scenario, the mosaic nature of the extant eukaryotic genomes resulted from the ‘inherited chimerism’ of the lone mitochondria bacterial ancestor because of the propensity of bacteria to participate in HGT with distantly related groups (Ku et al. 2015a, 2015b, Koonin 2015).

We suggest that the data we report here support the gradualist view of organelle evolution. In particular, we find that the majority of nutrient-related HGTs occurred prior to the divergence of the Phenacoccinae and Pseudococcinae (Figures 3 and 4), and thus before the establishment of the gammaproteobacterial symbionts. This means that the patchwork pattern we observe in the lysine and riboflavin pathways, where gene products are differentially retained across two or three genomes in different ways in different insect lineages, have changed numerous times in mealybugs. HTGs in the riboflavin pathway show particular stability in their inferred interactions with gammaproteobacteiral genes of different acquisitions and ages (Figure 3). Likewise, the HTGs of the lysine pathway remain stable in all lineages as the gammaproteobacterial symbionts of different inferred ages lost genes in response to genes present on both *Tremblaya* and insect genomes (Figure 3). Our results make it clear that HTGs can remain stable on host genomes for millions of years, even after the addition or replacement of symbionts that share pathways with these genes, and directly show how mosaic metabolic pathways can be built gene by gene as symbionts come and go over time. We note that our results are remarkably consistent with the ‘shopping bag’ hypothesis for the evolution of plastids (Larkum et al. 2007). This hypothesis argues that establishment of an endosymbiosis should be regarded as a continuous process involving a number of partners rather than a single event involving two partners. A series of transient and unstable symbioses can be attempted before a relatively stable relationship is fixed (and eventually can give rise to an organelle). Genes from these transient symbionts can be, however, transferred to the host genome and used to support the next symbiont (Larkum et al. 2007). Of course, our data do not rule inherited chimerism as a contributor to the taxonomic diversity of HGTs that support organelle function, as many bacterial genomes are, of course, taxonomically mosaic in nature due to HGT (Ku et al. 2015a).

### Symbiont supplementation and replacement to claw out of the rabbit hole

At the onset of nutritional symbiosis, a new organism comes on board and allows access to a previously inaccessible food source. Rapid adaptation and diversification can occur—the new symbiont adapts to the host, the host to the symbiont, and the entire symbiosis expands in the newly available ecological niche. But cases where a bacterial symbiont takes up stable residence in a host cell also seem to lead to irreversible degeneration and co-dependence between host and symbiont (Moran 1996, Andersson and Andersson 1999, Fares et al. 2002b), a situation recently described as the “symbiotic rabbit hole” (Bennett and Moran 2015). What HGT, symbiont supplementation, and symbiont replacement may offer is a way out—at least temporarily, perhaps permanently—of this degenerative ratchet.

But new symbionts provide not only evolutionary reinvigoration, but also ecological opportunity. It is interesting to note that the mealybug with one of the broadest host ranges is the species with the most recent gammaproteobacterial replacement, *P. longispinus. P. longispinus* is an important agricultural pest, and is known to feed on plants from 82 families [http://scalenet.info/catalogue/pseudococcus%20longispinus/]. It seems possible that the fresh symbionts with large genomes could provide novel functions unavailable in more degenerate symbionts, and thus propel in the symbioses into new niches.

## Acknowledgements

FH was funded by the Fulbright Commission and the Grant Agency of the University of South Bohemia (04-001/2014/P). JPM was funded by the National Science Foundation grants IOS-1256680 and IOS-1553529, NSF-EPSCoR Award NSF-IIA-1443108 to the Montana Institute on Ecosystems, and the National Aeronautics and Space Administration Astrobiology Institute award NNA15BB04A. We thank Genomics Core Facility at the University of Montana, DNA Sequencing Facility at the University of Utah, and the EMBL Genomics Core Facility in Heidelberg for excellent sequencing services.

## Data Deposition

The nine complete endosymbiont genomes, five draft assemblies of insect genomes, and raw data were deposited into the European Nucleotide Archive (ENA) under the following study numbers: *Maconellicoccus hirsutus*: PRJEB12066; *Ferrisia virgata*: PRJEB12067, *Pseudococcus longispinus*: PRJEB12068; *Paracoccus marginatus:* PRJEB12069; *Trionymus perrisii*: PRJEB12071. Unannotated draft genomes of two Enterobacteriaceae symbionts from *Pseudococcoccus longispinus* mealybugs and a B-supergroup *Wolbachia* strain sequenced from *Maconellicoccus hirsutus* mealybugs were deposited in Figshare under Digital Object Identifier numbers 10.6084/m9.figshare.2010393 and 10.6084/m9.figshare.2010390.

## References

Andersson, J. O., and S. G. Andersson. 1999. Insights into the evolutionary process of genome degradation. Current Opinion in Genetics & Development 9:664–671.

Bankevich, A., S. Nurk, D. Antipov, A. A. Gurevich, M. Dvorkin, A. S. Kulikov, V. M. Lesin, S. I. Nikolenko, S. Pham, A. D. Prjibelski, A. V. Pyshkin, A. V. Sirotkin, N. Vyahhi, G. Tesler, M. A. Alekseyev, and P. A. Pevzner. 2012. SPAdes: a new genome assembly algorithm and its applications to single-cell sequencing. Journal of Computational Biology 19:455–477.

Belda, E., A. Moya, S. Bentley, and F. J. Silva. 2010. Mobile genetic element proliferation and gene inactivation impact over the genome structure and metabolic capabilities of Sodalis glossinidius, the secondary endosymbiont of tsetse flies. BMC Genomics 11:449.

Bennett, G. M., and N. A. Moran. 2013. Small, smaller, smallest: the origins and evolution of ancient dual symbioses in a phloem-feeding insect. Genome Biology and Evolution 5:1675–1688.

Bennett, G. M., and N. a. Moran. 2015. Heritable symbiosis: The advantages and perils of an evolutionary rabbit hole. Proceedings of the National Academy of Sciences 112:10169–10176.

Boisvert, S., F. Laviolette, and J. Corbeil. 2010. Ray: simultaneous assembly of reads from a mix of high-throughput sequencing technologies. Journal of Computational Biology 17:1519–1533.

Bolger, A. M., M. Lohse, and B. Usadel. 2014. Trimmomatic: a flexible trimmer for Illumina sequence data. Bioinformatics (Oxford, England) 30:1–7.

Booth, A., and W. F. Doolittle. 2015. Eukaryogenesis, how special really? Proceedings of the National Academy of Sciences 112:10278–10285.

Burger, G., M. W. Gray, L. Forget, and B. F. Lang. 2013. Strikingly bacteria-like and gene-rich mitochondrial genomes throughout jakobid protists. Genome Biology and Evolution 5:418–438.

Campbell, M. A., J. T. Van Leuven, R. C. Meister, K. M. Carey, C. Simon, P. John, J. T. Van Leuven, R. C. Meister, K. M. Carey, C. Simon, and J. P. McCutcheon. 2015. Genome expansion via lineage splitting and genome reduction in the cicada endosymbiont Hodgkinia. Proceedings of the National Academy of Sciences 112:10192–10199.

Capella-Gutiérrez, S., J. M. Silla-Martínez, and T. Gabaldón. 2009. trimAl: a tool for automated alignment trimming in large-scale phylogenetic analyses. Bioinformatics (Oxford, England) 25:1972–1973.

Clayton, A. L., K. F. Oakeson, M. Gutin, A. Pontes, D. M. Dunn, A. C. von Niederhausern, R. B. Weiss, M. Fisher, and C. Dale. 2012. A novel human-infection-derived bacterium provides insights into the evolutionary origins of mutualistic insect-bacterial symbioses. Plos Genetics 8:e1002990.

Cooper, B. S., C. R. Burrus, C. Ji, M. W. Hahn, and K. L. Montooth. 2015. Similar efficacies of selection shape mitochondrial and nuclear genes in both Drosophila melanogaster and Homo sapiens. G3 (Bethesda, Md.) 5:2165–2176.

Dale, C., S. A. Young, D. T. Haydon, and S. C. Welburn. 2001. The insect endosymbiont Sodalis glossinidius utilizes a type III secretion system for cell invasion. Proceedings of the National Academy of Sciences of the United States of America 98:1883–1888.

Darling, A. E., B. Mau, and N. T. Perna. 2010. progressiveMauve: multiple genome alignment with gene gain, loss and rearrangement. Plos One 5:e11147.

Delmont, T. O., and A. M. Eren. 2016. Identifying contamination with advanced visualization and analysis practices: metagenomic approaches for eukaryotic genome assemblies. PeerJ PrePrints 4:e1695v1.

Denton, J. F., J. Lugo-Martinez, A. E. Tucker, D. R. Schrider, W. C. Warren, and M. W. Hahn. 2014. Extensive error in the number of genes inferred from draft genome assemblies. PLoS Computational Biology 10:e1003998.

von Dohlen, C. D., S. Kohler, S. T. Alsop, and W. R. McManus. 2001. Mealybug β-proteobacterial endosymbionts contain β-proteobacterial symbionts. Nature 412:433–436.

Douglas, A. E. 1989. Mycetocyte symbiosis in insects. Biological Reviews of the Cambridge Philosophical Society 64:409–434.

Duncan, R. P., F. Husnik, J. T. Van Leuven, D. G. Gilbert, L. M. Dávalos, J. P. McCutcheon, and A. C. C. Wilson. 2014. Dynamic recruitment of amino acid transporters to the insect/symbiont interface. Molecular Ecology 23:1608–23.

Embley, T. M., and W. Martin. 2006. Eukaryotic evolution, changes and challenges. Nature 440:623–630.

Ettema, T. J. G. 2016. Evolution: Mitochondria in the second act. Nature:doi:10.1038/nature16876.

Fares, M. A., E. Barrio, B. Sabater-Munoz, and A. Moya. 2002a. The evolution of the heat-shock protein GroEL from Buchnera, the primary endosymbiont of aphids, is governed by positive selection. Molecular Biology and Evolution 19:1162–1170.

Fares, M. A., M. X. Ruiz-Gonzalez, A. Moya, S. F. Elena, and E. Barrio. 2002b. GroEL buffers against deleterious mutations. Nature 417:398.

Frank, A. C., H. Amiri, and S. G. Andersson. 2002. Genome deterioration: loss of repeated sequences and accumulation of junk DNA. Genetica 115:1–12.

Gatehouse, L. N., P. Sutherland, S. A. Forgie, R. Kaji, and J. T. Christeller. 2012. Molecular and histological characterization of primary (betaproteobacteria) and secondary (gammaproteobacteria) endosymbionts of three mealybug species. Applied and Environmental Microbiology 78:1187–1197.

Gray, M. W. 2015. Mosaic nature of the mitochondrial proteome: Implications for the origin and evolution of mitochondria. Proceedings of the National Academy of Sciences 112:10133–10138.

Gray, M. W., and W. F. Doolittle. 1982. Has the endosymbiont hypothesis been proven? Microbiological Reviews 46:1–42.

Gruwell, M. E., N. B. Hardy, P. J. Gullan, and K. Dittmar. 2010. Evolutionary relationships among primary endosymbionts in the mealybug subfamily Phenacoccinae (Hemiptera: Coccoidea: Pseudococcidae). Applied and Environmental Microbiology 76:7521–7525.

Gurevich, A., V. Saveliev, N. Vyahhi, and G. Tesler. 2013. QUAST: quality assessment tool for genome assemblies. Bioinformatics (Oxford, England) 29:1072–1075.

Hosokawa, T., N. Kaiwa, Y. Matsuura, Y. Kikuchi, and T. Fukatsu. 2015. Infection prevalence of Sodalis symbionts among stinkbugs. Zoological Letters 1:5.

Huerta-Cepas, J., J. Dopazo, and T. Gabaldón. 2010. ETE: a python environment for tree exploration. BMC Bioinformatics 11:24.

Hunt, M., T. Kikuchi, M. Sanders, C. Newbold, M. Berriman, and T. D. Otto. 2013. REAPR: a universal tool for genome assembly evaluation. Genome Biology 14:R47.

Husnik, F., N. Nikoh, R. Koga, L. Ross, R. P. Duncan, M. Fujie, M. Tanaka, N. Satoh, D. Bachtrog, A. C. C. Wilson, C. D. von Dohlen, T. Fukatsu, and J. P. McCutcheon. 2013. Horizontal gene transfer from diverse bacteria to an insect genome enables a tripartite nested mealybug symbiosis. Cell 153:1567–1578.

International Aphid Genomics Consortium. 2010. Genome sequence of the pea aphid Acyrthosiphon pisum. PLoS Biology 8:e1000313.

Jones, P., D. Binns, H.-Y. Chang, M. Fraser, W. Li, C. McAnulla, H. McWilliam, J. Maslen, A. Mitchell, G. Nuka, S. Pesseat, A. F. Quinn, A. Sangrador-Vegas, M. Scheremetjew, S.-Y. Yong, R. Lopez, and S. Hunter. 2014. InterProScan 5: genome-scale protein function classification. Bioinformatics (Oxford, England) 30:1236–40.

Karp, P. D., S. M. Paley, M. Krummenacker, M. Latendresse, J. M. Dale, T. J. Lee, P. Kaipa, F. Gilham, A. Spaulding, L. Popescu, T. Altman, I. Paulsen, I. M. Keseler, and R. Caspi. 2010. Pathway Tools version 13.0: integrated software for pathway/genome informatics and systems biology. Briefings in Bioinformatics 11:40–79.

Katoh, K., and H. Toh. 2008. Recent developments in the MAFFT multiple sequence alignment program. Briefings in Bioinformatics 9:286–298.

Keeling, P. J., and J. M. Archibald. 2008. Organelle evolution: what’s in a name? Current Biology 18:R345–7.

Keeling, P. J., J. P. McCutcheon, and W. F. Doolittle. 2015. Symbiosis becoming permanent: Survival of the luckiest. Proceedings of the National Academy of Sciences 112:10101–10103.

Kobiałka, M., A. Michalik, M. Walczak, Ł. Junkiert, and T. Szklarzewicz. 2015. Sulcia symbiont of the leafhopper Macrosteles laevis (Ribaut, 1927) (Insecta, Hemiptera, Cicadellidae: Deltocephalinae) harbors Arsenophonus bacteria. Protoplasma:10.1007/s00709–015–0854.

Koga, R., G. M. Bennett, J. R. Cryan, and N. a Moran. 2013. Evolutionary replacement of obligate symbionts in an ancient and diverse insect lineage. Environmental Microbiology 15:2073–2081.

Koga, R., and N. a Moran. 2014. Swapping symbionts in spittlebugs: evolutionary replacement of a reduced genome symbiont. The ISME journal 8:1237–1246.

Kono, M., R. Koga, M. Shimada, and T. Fukatsu. 2008. Infection dynamics of coexisting Beta-and Gammaproteobacteria in the nested endosymbiotic system of mealybugs. Applied and Environmental Microbiology 74:4175–4184.

Konwar, K. M., N. W. Hanson, A. P. Pagé, and S. J. Hallam. 2013. MetaPathways: a modular pipeline for constructing pathway/genome databases from environmental sequence information. BMC Bioinformatics 14:202.

Koonin, E. V. 2015. Archaeal ancestors of eukaryotes: not so elusive any more. BMC Biology 13:84.

Koutsovoulos, G., S. Kumar, D. R. Laetsch, L. Stevens, J. Daub, C. Conlon, H. Maroon, F. Thomas, A. Aboobaker, and M. Blaxter. 2015. The genome of the tardigrade Hypsibius dujardini. bioRxiv:10.1101/033464.

Ku, C., S. Nelson-Sathi, M. Roettger, S. Garg, E. Hazkani-Covo, and W. F. Martin. 2015a. Endosymbiotic gene transfer from prokaryotic pangenomes: Inherited chimerism in eukaryotes. Proceedings of the National Academy of Sciences of the United States of America 112:10139–10146.

Ku, C., S. Nelson-Sathi, M. Roettger, F. L. Sousa, P. J. Lockhart, D. Bryant, E. Hazkani-Covo, J. O. McInerney, G. Landan, and W. F. Martin. 2015b. Endosymbiotic origin and differential loss of eukaryotic genes. Nature 524:427–432.

Kumar, S., M. Jones, G. Koutsovoulos, M. Clarke, and M. Blaxter. 2013. Blobology: exploring raw genome data for contaminants, symbionts and parasites using taxon-annotated GC-coverage plots. Frontiers in Genetics 4:237.

Kurland, C. G., and S. G. Andersson. 2000. Origin and evolution of the mitochondrial proteome. Microbiology and Molecular Biology Reviews 64:786–820.

Lamelas, A., M. J. Gosalbes, A. Manzano-Marín, J. Peretó, A. Moya, A. Latorre, A. Manzano-Marin, and J. Pereto. 2011. Serratia symbiotica from the aphid Cinara cedri: A missing link from facultative to obligate insect endosymbiont. PLoS Genetics 7:e1002357.

Larkum, A. W. D., P. J. Lockhart, and C. J. Howe. 2007. Shopping for plastids. Trends in Plant Science 12:189–195.

Lartillot, N., N. Rodrigue, D. Stubbs, and J. Richer. 2013. PhyloBayes MPI: phylogenetic reconstruction with infinite mixtures of profiles in a parallel environment. Systematic Biology 62:611–615.

Lefevre, C., H. Charles, A. Vallier, B. Delobel, B. Farrell, and A. Heddi. 2004. Endosymbiont phylogenesis in the Dryophthoridae weevils: evidence for bacterial replacement. Molecular Biology and Evolution 21:965–973.

Li, L., C. J. Stoeckert, and D. S. Roos. 2003. OrthoMCL: identification of ortholog groups for eukaryotic genomes. Genome Research 13:2178–2189.

Lomsadze, A., V. Ter-Hovhannisyan, Y. O. Chernoff, and M. Borodovsky. 2005. Gene identification in novel eukaryotic genomes by self-training algorithm. Nucleic Acids Research 33:6494–506.

López-Madrigal, S., A. Beltra, S. Resurreccion, A. Soto, A. Latorre, A. Moya, and R. Gil. 2014. Molecular evidence for ongoing complementarity and horizontal gene transfer in endosymbiotic systems of mealybugs. Frontiers in Microbiology 5.

Luan, J.-B., W. Chen, D. K. Hasegawa, A. M. Simmons, W. M. Wintermantel, K.-S. Ling, Z. Fei, S.-S. Liu, and A. E. Douglas. 2015. Metabolic coevolution in the bacterial symbiosis of whiteflies and related plant sap-feeding insects. Genome Biology and Evolution 7:2635–47.

Manzano-Marín, A., and A. Latorre. 2014. Settling down: The genome of Serratia symbiotica from the aphid Cinara tujafilina zooms in on the process of accommodation to a cooperative intracellular life. Genome Biology and Evolution 6:1683–1698.

Martin, W., and M. Müller. 1998. The hydrogen hypothesis for the first eukaryote. Nature 392:37–41.

McCutcheon, J. P., and C. D. Von Dohlen. 2011. An interdependent metabolic patchwork in the nested symbiosis of mealybugs. Current Biology 21:1366–1372.

McCutcheon, J. P., and P. J. Keeling. 2014. Endosymbiosis: protein targeting further erodes the organelle/symbiont distinction. Current Biology 24:R654–655.

McCutcheon, J. P., B. R. McDonald, and N. A. Moran. 2009. Convergent evolution of metabolic roles in bacterial co-symbionts of insects. Proceedings of the National Academy of Sciences of the United States of America 106:15394–15399.

McCutcheon, J. P., and N. A. Moran. 2007. Parallel genomic evolution and metabolic interdependence in an ancient symbiosis. Proceedings of the National Academy of Sciences of the United States of America 104:19392–19397.

McCutcheon, J. P., and N. A. Moran. 2010. Functional convergence in reduced genomes of bacterial symbionts spanning 200 million years of evolution. Genome Biology and Evolution 2:708–718.

McCutcheon, J. P., and N. a. Moran. 2011. Extreme genome reduction in symbiotic bacteria. Nature Reviews Microbiology 10:13–26.

Michalik, A., W. Jankowska, M. Kot, A. Gołas, and T. Szklarzewicz. 2014. Symbiosis in the green leafhopper, Cicadella viridis (Hemiptera, Cicadellidae). Association in statu nascendi? Arthropod Structure & Development 43:579–87.

Moran, N. A. 1996. Accelerated evolution and Muller’s rachet in endosymbiotic bacteria. Proceedings of the National Academy of Sciences of the United States of America 93:2873–2878.

Moran, N. A. 2002. Microbial minimalism: genome reduction in bacterial pathogens. Cell 108:583–586.

Moran, N. A., and G. M. Bennett. 2014. The tiniest tiny genomes. Annual Review of Microbiology 68:195–215.

Moran, N. A., J. P. McCutcheon, and A. Nakabachi. 2008. Genomics and evolution of heritable bacterial symbionts. Annual Review of Genetics 42:165–190.

Moran, N. A., and A. Mira. 2001. The process of genome shrinkage in the obligate symbiont Buchnera aphidicola. Genome Biology 2:54.

Moya, A., J. Pereto, R. Gil, A. Latorre, and J. Peretó. 2008. Learning how to live together: genomic insights into prokaryote-animal symbioses. Nature Reviews Genetics 9:218–229.

Nakabachi, A., K. Ishida, Y. Hongoh, M. Ohkuma, and S. Miyagishima. 2014. Aphid gene of bacterial origin encodes a protein transported to an obligate endosymbiont. Current Biology 24:R640–641.

Nakabachi, A., R. Ueoka, K. Oshima, R. Teta, A. Mangoni, M. Gurgui, N. J. Oldham, G. van Echten-Deckert, K. Okamura, K. Yamamoto, H. Inoue, M. Ohkuma, Y. Hongoh, S. Miyagishima, M. Hattori, J. Piel, and T. Fukatsu. 2013. Defensive bacteriome symbiont with a drastically reduced genome. Current Biology 23:1478–84.

Nakayama, T., and K. Ishida. 2009. Another acquisition of a primary photosynthetic organelle is underway in Paulinella chromatophora. Current Biology 19:R284–285.

Nikoh, N., J. P. McCutcheon, T. Kudo, S. Y. Miyagishima, N. A. Moran, and A. Nakabachi. 2010. Bacterial genes in the aphid genome: absence of functional gene transfer from Buchnera to its host. Plos Genetics 6:e1000827.

Nowack, E. C. M., and A. R. Grossman. 2012. Trafficking of protein into the recently established photosynthetic organelles of Paulinella chromatophora. Proceedings of the National Academy of Sciences of the United States of America 109:5340–5345.

Nowack, E. C. M., and M. Melkonian. 2010. Endosymbiotic associations within protists. Philosophical transactions of the Royal Society of London. Series B, Biological sciences 365:699–712.

Nowack, E. C. M., H. Vogel, M. Groth, A. R. Grossman, M. Melkonian, and G. Glöckner. 2011. Endosymbiotic gene transfer and transcriptional regulation of transferred genes in Paulinella chromatophora. Molecular Biology and Evolution 28:407–22.

Oakeson, K. F., R. Gil, A. L. Clayton, D. M. Dunn, A. C. von Niederhausern, C. Hamil, A. Aoyagi, B. Duval, A. Baca, F. J. Silva, A. Vallier, D. G. Jackson, A. Latorre, R. B. Weiss, A. Heddi, A. Moya, and C. Dale. 2014. Genome degeneration and adaptation in a nascent stage of symbiosis. Genome Biology and Evolution 6:76–93.

Ochoa de Alda, J. A. G., R. Esteban, M. L. Diago, and J. Houmard. 2014. The plastid ancestor originated among one of the major cyanobacterial lineages. Nature Communications 5:4937.

Palmer, J. D. 1997. Organelle genomes: going, going, gone! Science 275:790–791.

Parra, G., K. Bradnam, and I. Korf. 2007. CEGMA: a pipeline to accurately annotate core genes in eukaryotic genomes. Bioinformatics (Oxford, England) 23:1061–1067.

Pittis, A. A., and T. Gabaldón. 2016. Late acquisition of mitochondria by a host with chimaeric prokaryotic ancestry. Nature:10.1038/nature16941.

Popadin, K. Y., S. I. Nikolaev, T. Junier, M. Baranova, and S. E. Antonarakis. 2013. Purifying selection in mammalian mitochondrial protein-coding genes is highly effective and congruent with evolution of nuclear genes. Molecular Biology and Evolution 30:347–55.

Ronquist, F., and J. P. Huelsenbeck. 2003. MrBayes 3: Bayesian phylogenetic inference under mixed models. Bioinformatics 19:1572–1574.

Rosenblueth, M., L. Sayavedra, H. Sámano-Sánchez, A. Roth, and E. Martínez-Romero. 2012. Evolutionary relationships of flavobacterial and enterobacterial endosymbionts with their scale insect hosts (Hemiptera: Coccoidea). Journal of Evolutionary Biology 25:2357–68.

Ruby, J. G., P. Bellare, and J. L. Derisi. 2013. PRICE: software for the targeted assembly of components of (Meta) genomic sequence data. G3 (Bethesda, Md.) 3:865–880.

Rutherford, K., J. Parkhill, J. Crook, T. Horsnell, P. Rice, M. A. Rajandream, and B. Barrell. 2000. Artemis: sequence visualization and annotation. Bioinformatics (Oxford, England) 16:944–945.

Seemann, T. 2014. Prokka: rapid prokaryotic genome annotation. Bioinformatics (Oxford, England) 30:2068–9.

Segata, N., D. Börnigen, X. C. Morgan, and C. Huttenhower. 2013. PhyloPhlAn is a new method for improved phylogenetic and taxonomic placement of microbes. Nature Communications 4:2304.

Simão, F. A., R. M. Waterhouse, P. Ioannidis, E. V Kriventseva, and E. M. Zdobnov. 2015. BUSCO: assessing genome assembly and annotation completeness with single-copy orthologs. Bioinformatics (Oxford, England) 31:3210–3212.

Sloan, D. B., A. J. Alverson, J. P. Chuckalovcak, M. Wu, D. E. McCauley, J. D. Palmer, and D. R. Taylor. 2012. Rapid evolution of enormous, multichromosomal genomes in flowering plant mitochondria with exceptionally high mutation rates. PLoS Biology 10:e1001241.

Sloan, D. B., and N. A. Moran. 2012a. Endosymbiotic bacteria as a source of carotenoids in whiteflies. Biology Letters 8:986–989.

Sloan, D. B., and N. A. Moran. 2012b. Genome reduction and co-evolution between the primary and secondary bacterial symbionts of psyllids. Molecular Biology and Evolution 29:3781–3792.

Sloan, D. B., A. Nakabachi, S. Richards, J. Qu, S. C. Murali, R. a. Gibbs, and N. a. Moran. 2014. Parallel histories of horizontal gene transfer facilitated extreme reduction of endosymbiont genomes in sap-feeding insects. Molecular Biology and Evolution 31:857–871.

Smith, D. R., and P. J. Keeling. 2015. Mitochondrial and plastid genome architecture: Reoccurring themes, but significant differences at the extremes. Proceedings of the National Academy of Sciences 112:10177–10184.

Smith, W. A., K. F. Oakeson, K. P. Johnson, D. L. Reed, T. Carter, K. L. Smith, R. Koga, T. Fukatsu, D. H. Clayton, and C. Dale. 2013. Phylogenetic analysis of symbionts in feather-feeding lice of the genus Columbicola: evidence for repeated symbiont replacements. BMC Evolutionary Biology 13:109.

Stamatakis, A. 2014. RAxML version 8: a tool for phylogenetic analysis and post-analysis of large phylogenies. Bioinformatics (Oxford, England) 30:1312–1313.

Stewart, F. J., I. L. G. Newton, and C. M. Cavanaugh. 2005. Chemosynthetic endosymbioses: adaptations to oxic–anoxic interfaces. Trends in Microbiology 13:439–448.

Tamas, I., L. Klasson, B. Canbäck, a K. Näslund, A.-S. S. Eriksson, J. J. Wernegreen, J. P. Sandström, N. A. Moran, S. G. E. Andersson, B. Canback, A. K. Naslund, A.-S. S. Eriksson, J. J. Wernegreen, J. P. Sandstrom, N. A. Moran, S. G. E. Andersson, B. Canbäck, a K. Näslund, and J. P. Sandström. 2002. 50 million years of genomic stasis in endosymbiotic bacteria. Science 296:2376–2379.

Thao, M. L., M. A. Clark, L. Baumann, E. B. Brennan, N. A. Moran, and P. Baumann. 2000. Secondary endosymbionts of psyllids have been acquired multiple times. Current Microbiology 41:300–304.

Thao, M. L., P. J. Gullan, and P. Baumann. 2002. Secondary (gamma-Proteobacteria) endosymbionts infect the primary (beta-Proteobacteria) endosymbionts of mealybugs multiple times and coevolve with their hosts. Applied and Environmental Microbiology 68:3190–3197.

Theissen, U., and W. Martin. 2006. The difference between organelles and endosymbionts. Current Biology 16:1016–1017.

Timmis, J. N., M. A. Ayliffe, C. Y. Huang, and W. Martin. 2004. Endosymbiotic gene transfer: organelle genomes forge eukaryotic chromosomes. Nature Reviews Genetics 5:123–135.

Toh, H., B. L. Weiss, S. A. H. Perkin, A. Yamashita, K. Oshima, M. Hattori, and S. Aksoy. 2006. Massive genome erosion and functional adaptations provide insights into the symbiotic lifestyle of Sodalis glossinidius in the tsetse host. Genome Research 16:149–156.

Vogel, K. J., and N. A. Moran. 2013. Functional and evolutionary analysis of the genome of an obligate fungal symbiont. Genome Biology and Evolution 5:891–904.

Walker, B. J., T. Abeel, T. Shea, M. Priest, A. Abouelliel, S. Sakthikumar, C. A. Cuomo, Q. Zeng, J. Wortman, S. K. Young, and A. M. Earl. 2014. Pilon: an integrated tool for comprehensive microbial variant detection and genome assembly improvement. Plos One 9:e112963.

Wang, Z., and M. Wu. 2015. An integrated phylogenomic approach toward pinpointing the origin of mitochondria. Scientific Reports 5:7949.

Wheeler, T. J., and S. R. Eddy. 2013. nhmmer: DNA homology search with profile HMMs. Bioinformatics (Oxford, England) 29:2487–2489.

Zimorski, V., C. Ku, W. F. Martin, and S. B. Gould. 2014. Endosymbiotic theory for organelle origins. Current Opinion in Microbiology 22:38–48.

